# The ethics conundrum in Recall by Genotype (RbG) research: Perspectives from birth cohort participants

**DOI:** 10.1101/124636

**Authors:** Joel T Minion, Frances Butcher, Nicholas Timpson, Madeleine J Murtagh

**Affiliations:** Policy, Ethics and Life Sciences (PEALS) Research Centre, School of Geography, Politics and Sociology, Newcastle University, UK; Oxford School of Public Health, Oxford, UK; MRC Integrative Epidemiology Unit, School of Social and Community Medicine, University of Bristol, Bristol, UK

**Author notes:** Correspondence: Prof Madeleine Murtagh.

**Keywords:** Recall by Genotype (RbG), The Avon Longitudinal Study of Parents and Children (ALSPAC), Children of the 90s (Co90s), qualitative, ethics, return of results, solidarity, reciprocity, co-production

## Abstract

Recall by genotype (RbG) research involves recruiting participants on the basis of genetic variation. The recent use of this approach in the Avon Longitudinal Study of Parents and Children (ALSPAC) has presented an important challenge for ethical conduct: for example, to inform participants of their genetic information and to deviate from existing policies of non-disclosure of results and risk unanticipated harms, or mask the full structure of the study design and miss an opportunity to open a process of disclosure within genotype directed research. Here we report analysis of 53 semi-structured interviews conducted with young adult ALSPAC participants. We found that the deep trust and faith participants developed over their long-term relationship with the study, alongside a naturally limited knowledge of genetics and modest interest in reported research outcomes, meant most reported few immediate concerns about being recruited by genotype. Participants considered themselves part of the ALSPAC team and in this vein identified constructive concerns about being informed of RbG research in recruitment documents as well as what general results would be most valuable and informative. Our findings highlight the importance of solidarity, reciprocity and co-production in biobank/participant relations, especially in long-term birth cohort studies where relationships develop over a lifetime. We argue that strong trusting relationships between study and participant confer great responsibility on researchers regarding duty of care. We make recommendations for conducting RbG research in longitudinal studies beyond those already available in the literature for other study-types.

**Conflict of Interest:** The authors declare no conflict of interest.

## Introduction

Recall by genotype (RbG) studies recruit prospective participants on the basis of a specific genetic variant or variants of interest. This contrasts with more established genetic research methodologies such as genome wide association studies (GWAS), where participants are selected to be representative specific populations or on the basis of particular phenotypes.

The likelihood of participants having a genetic variant of interest depends upon the frequency of the given variant (or variants) or on the predictability of genotype from phenotype (a function of penetrance and genetic architecture). For the purposes of characterising a specific genetic variant or impact of a specific collection of variants on detailed and new phenotypes of interest, this GWAS approach can be an inefficient and potentially costly recruitment mechanism, especially for rare diseases variants or for low frequency variants in more common diseases. In contrast, the RbG research design targets individuals who have already undergone genotyping, thereby allowing for smaller, more precise and efficient studies.

At the same time RbG studies present a novel ethical context and challenges. If participants are to be fully informed of the structure of an RbG study, including naming the variant(s) of interest, then inadvertent disclosure of genetic information during the recruitment process can be possible. Revealing this information has the potential for heightened anxiety and worry among participants about their health status, especially if a variant is implicated elsewhere in serious or stigmatising conditions. Here the ethical principle of autonomy (the basis of informed consent in contemporary research) is pitched against the ‘right not to know’ and the principle of non-maleficence (the precept to do no harm)^1^. This central tension in RbG research (also termed Genotype Driven Recruitment (GDR)) has been identified^2-4^ in studies involving biobanks and other repositories that include participants with genotypic data collection, including disease and tissue-based biobanks^5-7^, population-based biobanks^5^ and collections based on health records or direct to consumer testing^8^.

While careful not to conflate the issues of return of clinically useful findings (incidental findings) with disclosure of genetic information through recruitment, these studies point to a range of cognate issues that bear consideration: the need to avoid participant anxiety if the genetic information – or re-contact itself – is unexpected^2,8^; the possibility that even uncertain information may be important to participants^7^; and the challenge that informational utility includes personal utility and personal meaning as well as clinical utility, especially where there is parental, familial or personal experience of disease^5-7,9^.

Moreover, empirical studies with participants of disease-based biobanks demonstrate participant solidarity^10^ with research and researchers, and perceived co-production^11^ of the outcomes of research. Participants have had long-term relationships with their health care providers and researchers (who may have been one in the same) and often felt part of the ‘team’ working towards understanding and treatments for themselves, their family and the community affected by the disease^5,7^. Conversely, population-based biobank participants were more likely to be anxious about the disclosure of unexpected genetic information^5^.

Seven broad recommendations for RbG/GDR were developed on the basis of research participant interviews (mostly disease-based biobank participants), a survey with IRB Chairs and a consensus workshop^12^. While useful and important, these recommendations are context specific and do not work for all circumstances^8^. As noted by Olson and colleagues, guidelines are needed for “all the varied circumstances under which genetic data may become available for researchers”^4^. To this end, we examined the practical and ethical conundrum facing RbG studies in the context of a longitudinal birth cohort, the Avon Longitudinal Study of Parents and Children (ALSPAC).

ALSPAC is a “transgenerational prospective observational study investigating influences on health and development across the life course” and includes repeated deep phenotypic data and biosample collection, with whole genome sequencing on a subset of the cohort^13^.

Participants comprise a cohort of offspring born to pregnant women recruited between April 1991 and December 1992 in Bristol, UK, their parents, grandparents, siblings and most recently their own offspring^13^. The birth cohort (now adults) are also referred to (and frequently refer to themselves) as the Children of the 90s (Co90s). Alongside their families, participants have been followed through a series of ongoing data collection time points involving questionnaires and clinical assessment. Additionally, sub-studies involving smaller samples continue to be conducted, including now RbG studies. In all sub-studies it is ALSPAC administrators who circulate study invitations to existing participants on behalf of research teams within and beyond ALSPAC. Such invites arrive on Co90s letterhead and are signed by the ALSPAC principal investigators. Measures are taken to ensure cohort members are not overburdened with study invitations. To conduct research involving cohort members, sub-study researchers first work with ALSPAC administrators to ensure all recruitment materials meet established standards and protocols. Following the principles and methodologies of Responsible Research and Innovation (RRI)^14,15^, ALSPAC members are engaged in the approval processes for all research conducted within ALSPAC and take part in governance decisions. Each study application is reviewed by the Original Cohort Advisory Panel (OCAP), a committee of ALSPAC participants who provide input and advice on study design, methodology and acceptability based on their expertise as participants. Study applications are then submitted to the ALSPAC Ethics and Law Committee (ALEC) for final approval. ALSPAC participants comprise half the membership of the ALEC, which is currently chaired by a parent participant and deputy chaired by one of the Original Cohort. It was previously chaired by one of the authors (MJM).

Genome-wide common single nucleotide polymorphism (SNP) data are now available for over 8000 cohort members, including a sub-sample of approximately 1800 members for whom there is complete genome sequencing^16^. Since 2010, RbG studies involving ALSPAC members and genetic variants associated with topics as diverse as sleep, schizophrenia, smoking behaviour and platelet function^17^. ALSPAC maintains a general principle that biomedical information is not disclosed to cohort members unless there is clear evidence the benefits outweighs the risks and that three conditions are met: (1) data provide clear, unequivocal evidence of an existing or future health problem; (2) said health problem is amenable to treatment of proven benefit; and (3) the participant has indicated in advance he/she wishes to be informed if such a problem is identified (www.bristol.ac.uk/media-library/sites/alspac/migrated/documents/alspac-disclosure-policy.pdf). In participant recruitment, whilst the method and nature of recruitment is fully described, documents used to date, the specific genetic variation(s) examined in these RbG studies have not been communicated to potential participants

With no available empirical evidence of the views of the wider ALSPAC participant population we, therefore, undertook a qualitative study of participant perspectives on the recruitment of cohort members into RbG research and the possible receipt of individual genetic information. Members of this cohort are well placed for RbG studies because they have already been recruited, it is possible to contact them and in many cases have already given consent for genetic data to be used for future research. Such characteristics, however, as well as members’ ongoing relationship with ALSPAC could leave them potentially vulnerable to over-recruitment or foster a default willingness to consent. Given such challenges, our study therefore sought both to contribute empirical data and analysis to the scientific literature and to improve how ALSPAC oversees the research it approves and undertakes.

## Methods

Ethical approval for the study was obtained from the ALEC (Ref: 13341).

A purposive sample (i.e. a non-probability sample of participants based on our research objectives) was generated to include three categories: (1) individuals from the general ALSPAC cohort who had never participated in an RbG study; (2) individuals who had participated in an ALSPAC RbG study; and (3) individuals who had served on one or more ALSPAC committees at which RbG study applications were discussed (e.g. ALEC, OCAP). In May 2016, 200 study invitations were mailed to ALSPAC participants (**Table 1**). Expression of interest forms were received from 74 individuals, who were then contacted by the study researchers (FB and JTM). Two additional forms were received too late to be included in the study. Semi-structured interviews were subsequently conducted with 53 participants in person, by phone or via Facetime/Skype. In keeping with established ALSPAC practice, participants received a £20 gift voucher for taking part and were reimbursed for travel costs (where applicable (e.g. bus or taxi fares). An interview topic guide was revised iteratively throughout the data collection process to pursue and refine various lines of inquiry, with five main topic areas covered across all interviews (**Table 2**). Feedback was also solicited during the interviews on a leaflet developed to explain the principles of RbG research^18^. Copies of the leaflet were distributed with all study invitations as well as provided electronically prior to each interview upon request. The interviews lasted between 14 and 62 minutes and were audio-recorded and transcribed verbatim with each participant’s consent. The resultant transcripts were then checked for accuracy and anonymised.

**Table 1:**
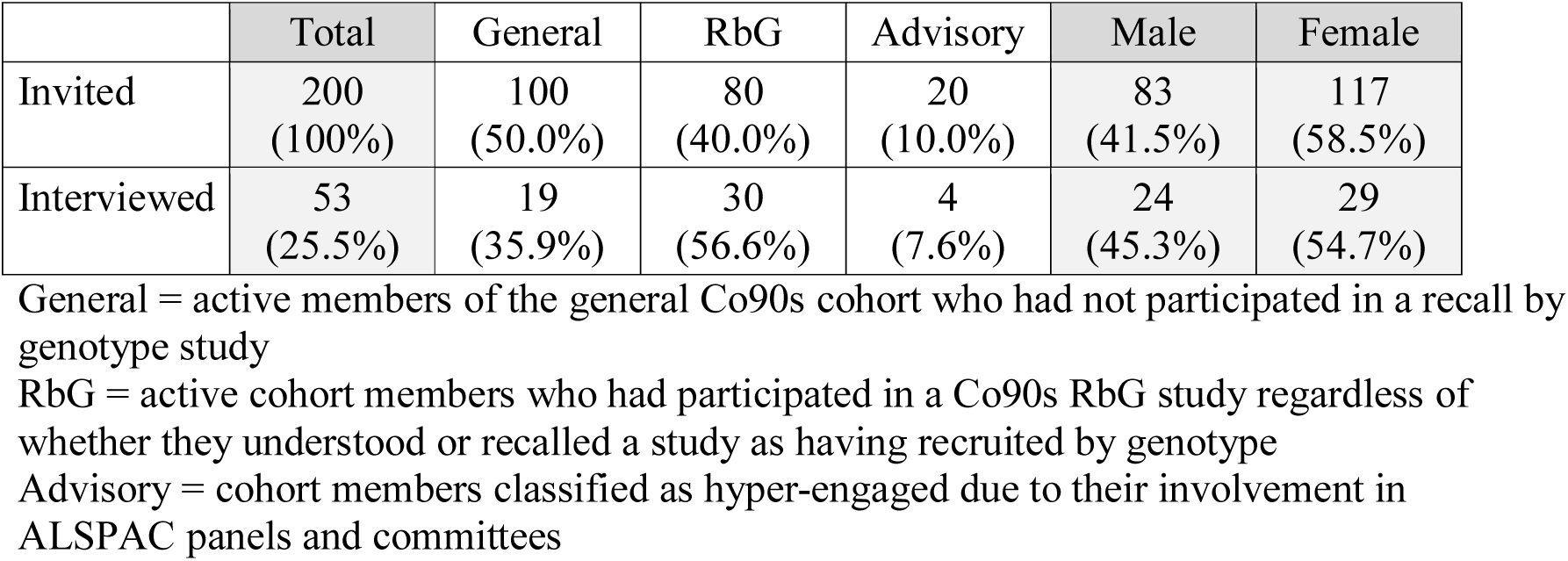
Participant recruitment.

|  | Total | General | RbG | Advisory | Male | Female |
| --- | --- | --- | --- | --- | --- | --- |
| Invited | 200<br>(100%) | 100<br>(50.0%) | 80<br>(40.0%) | 20<br>(10.0%) | 83<br>(41.5%) | 117<br>(58.5%) |
| Interviewed | 53<br>(25.5%) | 19<br>(35.9%) | 30<br>(56.6%) | 4<br>(7.6%) | 24<br>(45.3%) | 29<br>(54.7%) |

General = active members of the general Co90s cohort who had not participated in a recall by genotype study
RbG = active cohort members who had participated in a Co90s RbG study regardless of whether they understood or recalled a study as having recruited by genotype
Advisory = cohort members classified as hyper-engaged due to their involvement in ALSPAC panels and committees

**Table 2:** Topic areas discussed during interviews.

|  |  |
| --- | --- |
| Participating in ALSPAC research | Deciding whether to participate in any ALSPAC research when invited<br>Views on being told the reason for one's recruitment<br>Current knowledge and perspectives of RbG research<br>Feedback on the leaflet circulated explaining RbG research |
| Return of results | Perspectives on receiving general study findings (i.e. what did the study discover?)<br>Perspectives on receiving personal study results (including incidental or adverse findings) |
| Recall by genotype research | Thoughts on potentially being recruited into RbG studies (individuals were <u>not</u> informed whether they had already been involved in one)<br>Importance of knowing whether they had the specific genetic variation in question when recruited into a RbG study<br>Perspectives on receiving results (general or personal) from RbG studies versus non-RbG research<br>Thoughts on receiving results (general or personal) from a hypothetical RbG study linking a genetic variation to the increased probability of developing type 2 diabetes in middle age<br>Thoughts on the return of personal results from RbG research if doing so diverted funds from other research initiatives |
| General questions | General reflections on having been an ALSPAC cohort member<br>Perspectives on the importance of cohort member input into decisions about how ALSPAC operates and what research is approved<br>Ideas about what research topics ALSPAC might include or examine in greater detail in the future |
| Conclusion | Thoughts not otherwise captured during the interview<br>Ideas for additional interview questions or topics |

The interviews were analysed using the constant comparative method^19^. Two authors (JTM, MJM) familiarised themselves with the data by repeatedly and independently reading all of the transcripts and listening to the audio-recordings. A third author (FB), who had conducted 20 of the interviews for a Masters dissertation, worked similarly on those data under the direction of MJM. These results were then pooled with the main analysis. JTM and MJM analysed recurrent themes and emergent interpretations across several meetings, identifying open codes that were subsequently organised into a smaller number of concepts. JTM developed a first and second order coding frame and coded all transcripts using NVivo 11 software. MJM then undertook additional checks on coding and interpretation. Disconfirming cases (instances where participant perspectives ran counter to the developing interpretations) were sought throughout analysis to challenge emergent constructs, evaluate alternate explanations and refine resultant interpretations. Once written, the results were reviewed against the coded data one final time to ensure accuracy. In keeping with standard qualitative practice, brief excerpts of participants’ comments are provided in support of our analytic findings. All individuals quoted were pseudonymised specifically for this paper and all participant names are pseudonyms (nominated A-Z through the paper and indicative of gender only). The data for this study will be made available under managed access by ALSPAC at the end of the overarching qualitative study in 2018.

## Results and Discussion

53 participants were interviewed between June and August 2016 (**Table 1**), at which time theoretical saturation (the point at which no new themes emerged from the data) was judged (by MJM and JTM) to have been reached and recruitment ceased. 29 participants were female and 51 had been enrolled in ALSPAC continuously from birth (one participant reported having withdrawn only to re-enrol later, while another reported joining as a teenager). We include no further participant characteristics in order to maintain the privacy of the individuals involved. This paper reports three key perspectives identified in the data regarding how participants felt about recruitment and the return of research information more generally, and how these sentiments related to their perspectives on RbG research in particular.

### Perspectives on participation in research

As members of a birth cohort, all of the participants (except the two described earlier) had contributed data and biosamples continuously since the early 1990s. It was not surprising therefore that they reported very high levels of trust and engagement in both the administration and the scientific goals of the ALSPAC project. Participants frequently expressed a sense of satisfaction at having played an important role in the advancement of medical research and discovery of new treatments. When asked how they decided whether to accept new research invitations from ALSPAC, almost all of the participants responded that their default choice was to participate. Moreover, many reported that at times they gave only limited consideration to the research topic under consideration or the information provided in the recruitment documents.

> *Because I’ve been part of Children of the ‘90s all my life, I kind of just don’t really think about it. I just do it. It’s just part of something that I do now. So, yeah, I don’t tend to think, “Oh, should I do it or should I not do it?” I tend to just do whatever I can to help, really, because it’s such a big study and they’ve found out so much information that I think if I can help then, yeah, so I just do it*.
>
> — [Alice]

Few participants were able to provide examples of having declined a study invitation, whether RbG or any other study type. Where they could, only two reasons were given: practical barriers (e.g. lack of time; geographical distance) or personal dislikes (e.g. drawing blood; breast examinations).

Overall, participants expressed limited interest in knowing why they had been invited to a study. Several spoke about their past involvement in ALSPAC sub-studies using imprecise terms that suggested limited engagement with the research topic. (*I’ve taken part in, I think, pretty much every study that’s come up before. So I think there might have been some brain scan ones and a mammogram??* [Belinda]). While most participants did not feel it was important to know the reason for their recruitment, a few did indicate such information could be interesting. Almost none thought not knowing would deter their participation (*I’m not too bothered about why. It’s sometimes nice to know just out of interest, but it wouldn’t influence whether I said yes or no*. [Cathryn]). Despite limited interest in knowing the reasons for their recruitment, a small number of participants spoke about broader concerns regarding participation in research. They typically identified either ‘front end’ issues (e.g. gauging potential risk; agreeing with a study’s aims) or post-study concerns (e.g. potential return of unwelcome adverse findings).

> *I mean there is always that worry that, you know, something might be wrong with you and you might never find out even if all the data is collected. I mean I would – I guess I would be upset to find out later that I had some kind of terrible disease which I could have known about if I had been told why I was invited to something*.
>
> — [Deborah]

Such concerns, however, were almost never expressed as barriers to participation, nor did participants indicate any perceived need to discuss such matters with family, friends or ALSPAC staff.

The interview data also indicated that many participants used their ongoing trust and faith in ALSPAC to ‘shorthand’ recruitment decision-making. Participants frequently blurred any distinction between ALSPAC data collection events (commonly referred to as Focus Days) and sub-studies conducted by outside researchers. This finding did not lead us to conclude that participants were unable or unwilling to think critically about recruitment and participation; rather, we found most simply did not perceive a need to do so. The exception were participants who had served on ALSPAC panels and committees. While equally expressing high levels of trust and faith in ALSPAC, these individuals appeared to engage more thoroughly with study invitations, even if the outcome of whether to participate was similar to other participants.

### Perspectives on genetics

While our study did not specifically seek to gauge participants’ knowledge of genetics, the data suggested most had a basic high school level of understanding, with a smaller number either quite knowledgeable or largely uninformed.

Participants generally found the level of explanation in the RbG leaflet, which had been written at high school level knowledge, accessible for themselves and thought it would likely be so for other cohort members. Where participants reflected specifically on genetics in their interviews, they typically used terminology expressing curiosity or interest (*I’d just love to know really what’s different about me and how that maybe affects me compared to other people*. [Ella]) or perceived certainty (*Your DNA is essentially what you are built of. There is no escaping your own DNA*. [Fred]).

Given the iterative nature of the interview process, only the final 14 participants were asked specifically how much they thought about their genetic make-up in everyday life. Few reported doing so to any extent. Instead, personal genetics was expressed more as a “*backburner thing*” [George] of limited interest. In itself, this finding was not surprising given that the participants’ age and general circumstance (i.e. individuals in their mid-20s still establishing themselves in careers and personal relationships) would not otherwise suggest any pressing need to reflect regularly on their genetics. However, the perspectives of one participant – a new parent – did hint that the *status quo* could change significantly in the near future.

> *To be honest, I never used to really think about [my genetic make-up]. However, I do think about it more now. It’s something that, obviously with my little one, it’s something that you do think about and you do kind of wonder how it all works and things like that. So, yeah, it’s definitely something I think about more and more*.
>
> — [Alice]

With respect to genetics in relation to RbG research, almost none of the participants (besides those who had served on ALSPAC panels and committees) indicated much knowledge of this newer approach to recruitment. Given that over half our sample involved individuals who had already participated in an ALSPAC RbG study, this finding illustrated the known tendency of research participants to forget the details of research in which they are involved, specifically the information in consent forms and participant information materials (Lecouturier, 2012) but nonetheless must lead us to wonder whether recruitment information used in the past had been insufficiently clear or if participants simply had not engaged with this topic. When discussed in the interviews, RbG research was variously seen in either positive or neutral terms (“a *lot more efficient*” [Heather]; **“***just another study*” [Isabel]). Few participants felt they would be influenced in any way by knowing that they were being recruited to a study based on a genetic variation. Those. T who thought they would be influenced stated that such information might make them more rather than less likely to enrol.

The authors caution against interpreting this finding as evidence of unconditional acceptance of RbG research. It was not uncommon for our participants to weave subtle sentiments of unease or hesitation into their observations of genetics-based research. For example, when asked to reflect on possible disadvantages of RbG research, one participant observed:

> *I suppose [RbG studies] might reveal things that people don’t want to know, including negative outcomes. I suppose people don’t want to know that*.
>
> — [Joanne]

Another participant, one who knew she had taken part in an RbG study, expressed her concerns quite clearly:

> *I still think about it now because I still think – I guess I would have liked – I’m someone who would have liked to have known which group [I was in]. But there’s no way of telling me because then it would affect the study if I knew or it could affect it. But I still think – that was a year or two ago – and I still think, “Oh, I wonder which group I was part of?” because I wasn’t actually told*.
>
> — [Lacey]

While participants reported being largely unconcerned about RbG research generally, experiences such as Lacey’s could increase as RbG recruitment becomes more common and as cohort members have greater reason to reflect critically about being recruited based on genotype (e.g. once they have or are planning to have children themselves). Individuals who had seen little reason to question their possible involvement in an RbG study might possibly develop a stronger interest in doing so.

### Perspectives on the return research information

Our findings revealed an almost universal awareness and acceptance of ALSPAC’s practice to return biomedical information to cohort members only under very specific circumstances (REF). Almost everyone interviewed expressed little expectation they would receive information about themselves. Such data led us to conclude that ALSPAC’s non-disclosure policy was widely accepted for three reasons: it had been communicated repeatedly and clearly; individuals trusted ALSPAC to act if significant and remediable results were found; and participants viewed their role in ALSPAC primarily as data providers. As for the communication of research findings more generally, most participants reported being only marginally interested in learning the outcomes of ALSPAC studies and were happy to receive such information via newsletters on an occasional basis. When presented with the possibility of using technology to ‘push’ personalised digests to individuals on findings of specific interest to them (e.g. asthma), few saw much appeal. Similarly, there was little interest beyond curiosity in receiving routine individual results (despite many participants recalling fondly having received souvenir copies of scans and such as children).

While our participants did not generally express any strong concerns about the return of research information, the authors did identify two matters of note. First, ALSPAC was very much expected to observe its current policy. Participants articulated this expectation either as an ethical obligation or as a benefit arising from being a cohort member. This was a general expectation and not restricted to RbG studies or genetic information.

> *I went to my last Focus group and then I got a phone call from the [ALSPAC] doctor a week later because my blood results weren’t quite right. And they advised me to go to the doctor’s and have a repeat. And it was all fine in the end, but I think that was really good of them to call me – to let me know – because if I hadn’t have gone to Children of the 90s and hadn’t had that blood test, then it might have been something more serious and then I would have never have known about it*.
>
> — [Lacey]

The second concern noted among participants about the return of information related specifically to RbG research. During the interviews individuals were presented with a hypothetical scenario in which RbG research identified a link between a genetic variation and an increased likelihood of developing type 2 diabetes in middle age. Most participants felt ALSPAC had a duty to tell them both the outcome of such a study and whether they personally carried the variation in question (either at time of recruitment or following the study). The reasons cited turned less on ethical obligation or participatory benefit and more on a belief that health threats in the future might potentially be mitigated through behaviour change.

> *To be honest, I don’t know much about… type 2 diabetes, so if it was possible for me to prevent it by controlling my external factors then, yes, they should let me know. I mean, I think either way, yes, they should let me know. But if there was a way to prevent it, then it’s more important for them to let me know so I could do something about it and I’d have more control over it*.
>
> — [Katy]

Finally, participants were asked to consider the relative cost of returning individual results, especially given an anticipated increase in RbG research. While individuals expected ALSPAC to return adverse findings of clinical significance (which could include findings such as the hypothetical study on type 2 diabetes), most preferred funds be spent on research if the mechanics of returning results became too costly.

> *… in an ideal world we’d get both, so without having to take any budget or resources from the study or publications of research and be able to bring in counsellors or whatever, and be able to relay that information back on to me, that would be the ideal world. But if it’s not the case and it’s not feasible – if it’s not financially viable – then I wouldn’t have any gripes, I suppose, it wouldn’t stop me from wanting to do the study*.
>
> — [Jack]

This finding suggested that participants’ perspectives on the return of results could become more complex and dynamic as the number of RbG studies increases. Such research might challenge current ALSPAC policy on returning results if participants situate the health consequences of such findings far enough into the future that individuals feel they can effect change and prevent negative outcomes. In such a situation, cohort members may no longer be willing to prioritise research over returning results.

### Concluding remarks

A key issue in the undertaking of RbG research is the conflation of incidental findings (i.e. those that are a by-product of volunteering as a healthy participant for an otherwise hypothesis-free genetic study)^16^ with the screening of genetic variation in a study of otherwise clinically uninformative phenotypes. Where the variant (or variants) is associated with a serious or stigmatising condition such that an enterprising or curious participant can ‘google’ the variant names, a decision needs to be made about whether it is in the best interests of all potential participants to conceal that variant name. Importantly, this should not be done without the input of participants themselves, either through their routine involvement in study governance or through empirical studies such as this one. Such input must also not be deemed sufficient for all time; participants’ perspectives and needs are dynamic and will evolve. Indeed, the main concerns for RbG design are not the likely presence of incidental findings but the open and honest communication of study design in an effective manner to potential participants, and open discussion about the value of individual genotype status disclosure to participants and researchers given the chance of bias and inappropriate disclosure. In a study with strong participant engagement, such participants must be engaged fully in the determination of what counts as open communication and the value of individual genotype disclosure.

Our study demonstrated that participants’ perspectives on RbG research could only be fully understood within the context of such individuals’ long term and ongoing experience as participants in ALSPAC. Moreover, the context of individuals’ lives – including their involvement in genetic research – was dynamic and likely to evolve over time. In the case of ALSPAC, capturing perspectives on RbG research required acknowledging the expertise participants had acquired as data contributors over 20+ years and the role their relationship with ALSPAC played in their engagement with research. Like Michie’s^6^ Cystic Fibrosis patient, participants, ALSPAC participants’ demonstrated societal solidarity^10^, seeing themselves as contributing to a greater good. They also understood this contribution as reciprocal: participants appreciated learning something about themselves and expected ASLPAC to act in their interests as the need arose. Our results underscore the importance of learning from the experience of biobank participants when addressing the emergent governance challenges posed by RbG research and other new methodologies.

Although participants in our study seemingly expressed limited interest in knowing the reasons for their recruitment or the findings of such research, their attitudes speak to the ongoing advantages and limitations of the close working relationship between ALSPAC and its members. Indeed, the strength of trust in ALSPAC’s ‘duty of care’ role provides an important understanding, namely that any such perception or expectation arguably confers on the study a higher responsibility towards its participants than in a study where engagement is less strong.

While the ethical issues facing RbG research are closer to those considered in the context of population-based genetic research rather than family studies^20^, such an approach is inextricably linked to the issue of disclosing individual results^2^. There is undoubtedly a distinction between genetic data of clinical utility and data commonly used in epidemiological research, but differences in the implications of such categories for informed consent and ethical study conduct are often blurred^21^. We have attempted to contextualise for birth cohorts the “central tension” between avoiding the possibility of participant harm by revealing unwanted or misunderstood information and avoiding deception when explaining recruitment into RbG studies^12^. A limited number of studies have attempted to assess the effects of incorporating genetic information in intervention designs^22,23^ and some demonstrate the strong social and cultural determinants of those influences in the face of information about genetic susceptibility^24^. Despite this, evidence on the impact of using genetic data within recall studies is lacking.

### Recommendations

The seven recommendations that emerged from the consensus workshop about recruitment to studies based on genotype^12^ provide a good starting point for other biobanks. All have been followed by ALSPAC: (1) participants be made aware of potential for re-contact; (2) participants have a choice about whether to be re-contacted; (3) re-contact be made by a known person or entity; (4) recruitment be based on the biobank’s own processes; (5) thresholds for the return incidental findings be considered differently to the return of genetic information during recruitment; (6) genetic research information offered in the context of RbG recruitment should not leave participants uninformed about the study’s purpose; and (7) approaches to RbG recruitment be determined in consultation with ethics committees^12^. In the case of No.6, this must be tempered by the nature of the condition with which the variant is associated. Studies like ALSPAC that have a strong engagement ethic should build upon it by engaging not only researchers and ethics committees but also participants, who hold a separate and unique form of expertise. In fact, in the case of ALSPAC, it can draw not only its ethics committee but its participants’ panels and qualitative studies such as this. Between them, innovative approaches can be developed to provide information to participants in formats offering multiple levels of explanation; to advance creative mechanisms for communicating information on increasingly complex topics; and to offer opportunities for participants themselves to explore and critique the ethical implications of new study designs. Such biobanks must be mindful, however, in the context of a strong and established study-participant relationship, of the inclination of participants to decide first and ask questions later when considering study invitations. Attention to such understandings will underpin governance practices and help ensure they remain fit for purpose.

### Study limitations

While our findings may be suggestive of perspectives found among other birth cohorts, they are not likely transferable directly to other types of biobanks. Our sample was self-selected and as such we may have heard disproportionately from individuals interested in the topic and/or open to discussing a potentially sensitive subject matter. Nonetheless, this empirical study demonstrates that doing so is necessary to understand participants’ perspectives in terms of their wider experience of involvement in a biobank.

## Acknowledgements

We are extremely grateful to all the families who took part in this study, the midwives for their help in recruiting them, and the whole ALSPAC team, which includes interviewers, computer and laboratory technicians, clerical workers, research scientists, volunteers, managers, receptionists and nurses. The UK Medical Research Council and Wellcome (Grant ref: 102215/2/13/2) and the University of Bristol provide core support for ALSPAC. This publication is the work of the authors and MM will serve as guarantor for the contents of this paper. This research was specifically funded by the UK Medical Research Council, MRC Integrative Epidemiology Unit at the University of Bristol (MC_UU_12013/3).

